# From a biological template model to gait assistance with an exosuit

**DOI:** 10.1101/2020.03.24.005389

**Authors:** Vahid Firouzi, Ayoob Davoodi, Fariba Bahrami, Maziar A. Sharbafi

## Abstract

By invention of soft wearable assistive devices, known as exosuits, a new aspect in assisting unimpaired subjects is introduced. In this study, we designed and developed an exosuit with compliant biarticular thigh actuators, called BAExo. Unlike common method of using rigid actuators in exosuits, the BAExo is made of serial elastic actuators (SEA) resembling artificial muscles (AM). This bioinsipred design is complemented by the novel control concept of using the ground reaction force to adjust these AMs’ stiffness in the stance phase. By locking the motors in the swing phase the SEAs will be simplified to passive biarticular springs, which is sufficient for leg swinging. The key concept in our design and control approach is synthesizing human locomotion to develop assistive device, instead of copying the outputs of human motor control. Analysing human walking assistance using an experiment-based OpenSim model demonstrates the advantages of the proposed design and control of BAExo, regarding metabolic cost reduction and efficiency of the system. In addition, pilot experiments with the recently developed BAExo hardware support the applicability of the introduced method.

**Author summary:** Aging and mobility of elderly people are of crucial concern in developed countries. The U.S. Census Bureau reports that by the middle of the 21st century, about 80 million Americans will be 65 or older. According to the group’s research, medical costs resulting from falls by the elderly are expected to approach $32.4 billion by 2020. Therefore, assistance of elderly people and making the assistive devices more intelligent is a need in near future. However, this is not the only application of assistive devices. Exosuits, as soft wearable robots, introduced a new aspect in assisting a large range of population, even healthy young people. We introduce a novel design and control method for a new exosuit. As the research in the field of wearable assistive devices is growing in recent years and its application in daily life becomes more evident for the society, such studies with a unique view in design and control could have a significant impact. Our proposed biologically inspired approach could be potentially applied to other exosuits.

## Introduction

Lower limb exoskeletons for assisting people with decreased locomotion abilities has attracted the attention of researchers [1] and the healthcare industry [2, 3]. Assistive devices are also helpful for other categories of potential users, like firefighters, laborers and soldiers [4, 5]. They reduce the risk of injuries during work and improve the ergonomics of the work conditions. Also, recent studies show a huge societal concern about aging. For example, by aging, muscle force and power decrease and to reduce the cost of elderly and patients locomotion, assistive devices are demanded [6, 7].

**Table 1.** List of symbols, terms and definitions (*x* denotes one of the FMC, ABC or *P*)

| Nomenclature |  | Model Parameter |  |
| --- | --- | --- | --- |
| Abbreviation | Definition | Parameter | Definition |
| ABC | Adaptable Biarticular Compliance | $\tau_{VPP}$ | hip required torque in VPP model |
| BAExo | Bi-Articular Exosuit | $f_l$ | leg force |
| CMC | Computed Muscle Control | $l$ | leg length |
| CoM | Center of Mass | $r_h$ | distance from VPP to hip joint |
| DD | Direct Drive | $\psi$ | hip angle |
| EMG | Electromyography | $r_{VPP}$ | distance from CoM to VPP |
| FMC | Force Modulated Compliance | $\gamma$ | angle between body axis and the vector from CoM to VPP |
| FMCH | Force Modulated Compliant Hip | $\tau_h$ | hip torque in FMCH model |
| GRF | Ground Reaction Force | $c_h$ | normalized stiffness of the adjustable hip spring in the FMCH model |
| HAM | Hamstring | $\psi_0$ | rest angle of the adjustable hip spring in the FMCH model |
| PFF | Positive Force Feedback | $f_x$ | force applied by the actuator |
| RF | Rectus Femoris | $c$ | normalized stiffness in the FMC |
| SEA | Serial Elastic Actuator | $l_{AM}$ | length of the artificial muscle |
| SLIP | Spring Loaded Inverted Pendulum | $k$ | spring stiffness |
| TSLIP | Trunk + SLIP | $l_0^P$ | rest length of the passive spring mode |
| VPP | Virtual Pivot Point | $t_{lock}$ | motor's lock time |
| | | $l_0^F$ | reset length of the FMC mode |
| | | $T_f$ | FMC operation time interval |
| | | $T_p$ | passive spring operation time interval |
| | | $J$ | CMC's cost function |
| | | $a$ | muscles activation |
| | | $w$ | weight set |
| | | $J_A$ | CMC's assisted cost function |

One of the common goals of assistive device designers is to reduce the metabolic cost of locomotion [8]. This feature is important in all above-mentioned cases and situations. In the last couple of years researchers have succeeded in significantly improving the metabolic cost reduction in human gaits using powered exoskeletons [1, 9–11].

Powered exoskeletons can be categorized as (1) traditional ones with rigid structures which are connected to the body in parallel to the biological skeletal system (limb) [12], and (2) recently introduced exosuits using soft materials such as textiles [13, 14]. The ability to produce high torques, stabilize gaits in people with severe injuries (e.g., spinal cord) and apply such systems in rehabilitation are of the main advantages of the first group [15, 16]. In comparison, having light weight and low inertia, portability and the bioinspired design are of important features of the wearable exosuits. This later property results in more comfort and less constraints in movement by resolving misalignment between the subject’s and robot’s joints and the joints of the exoskeleton [13, 17]. Therefore, one can argue that soft exosuits are more appropriate than the rigid exoskeletons to be used by unimpaired human subjects for daily activities.

Because of tight interaction between humans and robots, control of the assistive devices becomes very crucial. Control strategies can be divided into model based and model-free approaches. Model-free control methods include (but not limited to^1^) time-based (predefined trajectory tracking based) approaches [20–23], predefined gait-pattern based control [24, 25], and EMG-based control [26]. These so-called black box methods [27] will have a high variance in walking economy among participants for fixed control strategies [11]. This problem can be solved by using a human-in-the-loop-optimization (HILO) [11, 28]. However, HILO is a time consuming and difficult process to optimize the accurate parameters for each individual.

Furthermore, it is not guaranteed that optimal parameters for one condition (such as a certain speed) will be an optimal parameter for all conditions (other speeds). On the other hand, issues exist in EMG-based control methods due to non-stationarity and subject-specificity [29]. Also in some cases, muscles are so close to each other that makes measurement of the surface EMG signal very difficult [29].

In contrary to black box methods, which use human locomotion control output signals (either joint torque/position or EMG signals) to control the assistive device, the white box (model-based) approaches try to understand the underlying control principles of human locomotion [30, 31]. However, dynamical systems (e.g., neuromuscular models [32]) that can describe human locomotion are complicated. The complexity of bioinspired model-based control of assistive devices can be reduced by model abstraction using the *Template & Anchor* concept [33], and by employing bioinspired morphological designs such as biarticular actuators [34, 35]. In short, we propose using human locomotion as a template for design and control of assistive devices (here, exosuit) instead of copying specific movement patterns (as in black box methods). Potentially, similarity in design and control of human and the robot locomotor systems can provide more synergistic behavior. A brief overview of the potential advantages of these lines of thoughts in the literature is presented in the following.

### Template-based control concept of GRF feedback

In [36], the force modulated hip compliance (FMCH model) was introduced as a template for posture control in which the GRF (ground reaction force) was used to tune the hip joint compliance. This method was later implemented in the LOPES II exoskeleton to assist human walking [37, 38]. The results revealed the advantages of using the force feedback in reducing the human metabolic cost of walking. From biological point of view, studies on normal and pathological gait demonstrated that humans also use proprioceptive feedback signals (e.g., detecting load as a complex parameter, recorded by very different types of receptors) in locomotion control [39, 40]. In addition to ability of measuring the load (e.g., body weight) in humans, high dependency of compensatory leg muscle activation was demonstrated experimentally [41].

### Facilitating control with compliant biarticular actuators

Studies on biomechanics of human gaits and robotics show that biarticular muscles help to generate motions in a more energy efficient way [34, 42]. In fact, these muscles actuate two joints simultaneously, transfer the energy towards distal joints and further control the output force direction [34, 43], [35]. Given these features, biarticular assistive devices can be more effective in reducing energy consumption during walking. Several examples for successful applications of the biarticular elements in designing assistive devices can be found in the literature [10, 37, 44, 45]. Different aspects of analyzing biarticular muscles from template modeling and biological locomotor systems to legged robot and assistive devices are reviewed in [35]. Moreover, adding compliance in a serial elastic actuation mechanism provides further advantages against the common method of using direct-drive motors for exosuit. Some of these advantages are (not limited to) recoiling energy, increasing robustness (e.g., at impacts), reducing peak power and required torques of the electric motors. In biarticular compliant actuators, the returned energy of one joint can be stored in the elastic element in part of the gait and returned in another joint in the following phase of gait [46, 47].

In this study, we introduce a new biarticular exosuit called *BAExo* that actuates hip and knee joints simultaneously. We investigate a **white box model-based control** in a novel actuator design with **compliant biarticular** actuation which was not employed in exosuits before. Our bioinspired white-box model is a simplified version of the reflex control [48] and is originated from the **GRF-based control**, supported by biomechanical studies [41, 49]. For overcoming control challenges we use 1) the aforementioned bioinspired control method of FMCH [36], in which the stiffness of the biarticular artificial muscles are tuned based on the GRF signal in the stance phase and 2) passive biarticular springs in the swing phase. Here, the advantages of using such a design and control of an exosuit are shown by analyzing human walking experimental data with an OpenSim model. The main advantages of the proposed design and control are demonstrated based on these experiment-based modeling outcomes. In addition, we performed pilot experiments on our recently developed *BAExo* hardware setup. The preliminary out comes support the applicability of the proposed method in real world. Further comprehensive experiments with more subjects and physiological signals (e.g., metabolics and EMG) measurement will be the future step.

## Materials and methods

This section describes the FMCH-based control approach, followed by details about data collection of the walking experiment and implementation of our gait assistance method with the biarticular exosuit (*BAExo*) in the OpenSim software. Finally, we explain implementation of the bioinspired ABC (Adaptable biarticular Compliance) control approach in the newly developed *BAExo* hardware.

### Control approach

Our control approach stems from template-based modeling of posture control in human locomotion. Therefore, first, we describe the VPP (virtual pivot point) concept [50] as a backbone of the FMCH model [36]. Then, we explain how to use the FMCH template model for control of exosuit in the stance phase (when the leg is in contact with the ground). For extension of this model for the segmented leg we use biarticular actuators with adjustable stiffness [51]. Then, control of the exosuit in the swing phase of walking (when the leg is moving freely in the air to take a step) will be described. The control scheme for the swing phase is also based on (passive) biarticular compliance, introduced in [52]. As both of these control strategies use biarticular thigh muscles, we can easily switch between them when the gait phase is changing (from stance to swing and vice versa). This hybrid design and control approach will be called as ABC (Adaptable biarticular Compliance), hereafter.

Legged locomotion can be described by three fundamental locomotor subfunctions [53]: 1) *Stance:* the axial function of the stance leg, 2) *Swing:* rotational movement of the swing leg and 3) *Balance:* posture control of the upper body. Here, we focus on the last two subfunctions for design and control of our exosuit. This is because the stance subfunction is mainly well (efficiently) supported by the human body in walking at normal speed.

#### Stance phase control

In order to describe our ABC method, first we explain the basic concept and developed control method for the balance locomotor subfunction which is based on the VPP concept, developed by Maus et al. [50].

##### The VPP model

The VPP which is observed in human and animal gaits is a point on the upper body above the CoM (center of mass) at which the ground reaction forces are intersecting during the stance phase [50]. This concept was already applied to predict hip torque required for posture control using template modelling for different gaits [54–56]. Template models are simple conceptual models using basic mechanical elements (e.g., mass, spring) which can explain some important features of locomotion [33]. One of the common templates for modeling walking and running is the SLIP (spring-loaded inverted pendulum) model [57–59]. This model consists of a point mass describing CoM, attached on top of a massless spring which represents the stance leg. For describing posture control, a rigid trunk can be added to the SLIP model resulting in the TSLIP (Trunk+SLIP) model [54–56]. Using this model, the required hip torque *τ*_*V PP*_ (see Fig. 1A) to redirect the GRF to a determined VPP can be calculated by

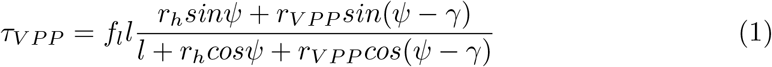

in which, *f*_*l*_, *l*, *ψ*, *r*_*V PP*_ and *r*_*h*_ are the leg force, leg length, hip angle, the distance from CoM to VPP and from VPP to hip joint, respectively. The VPP angle is defined by *γ*, the angle between body axis and the vector from CoM to VPP, as shown in Fig. 1A. For more details about derivation of VPP formulation please see [50].

**Fig 1.**
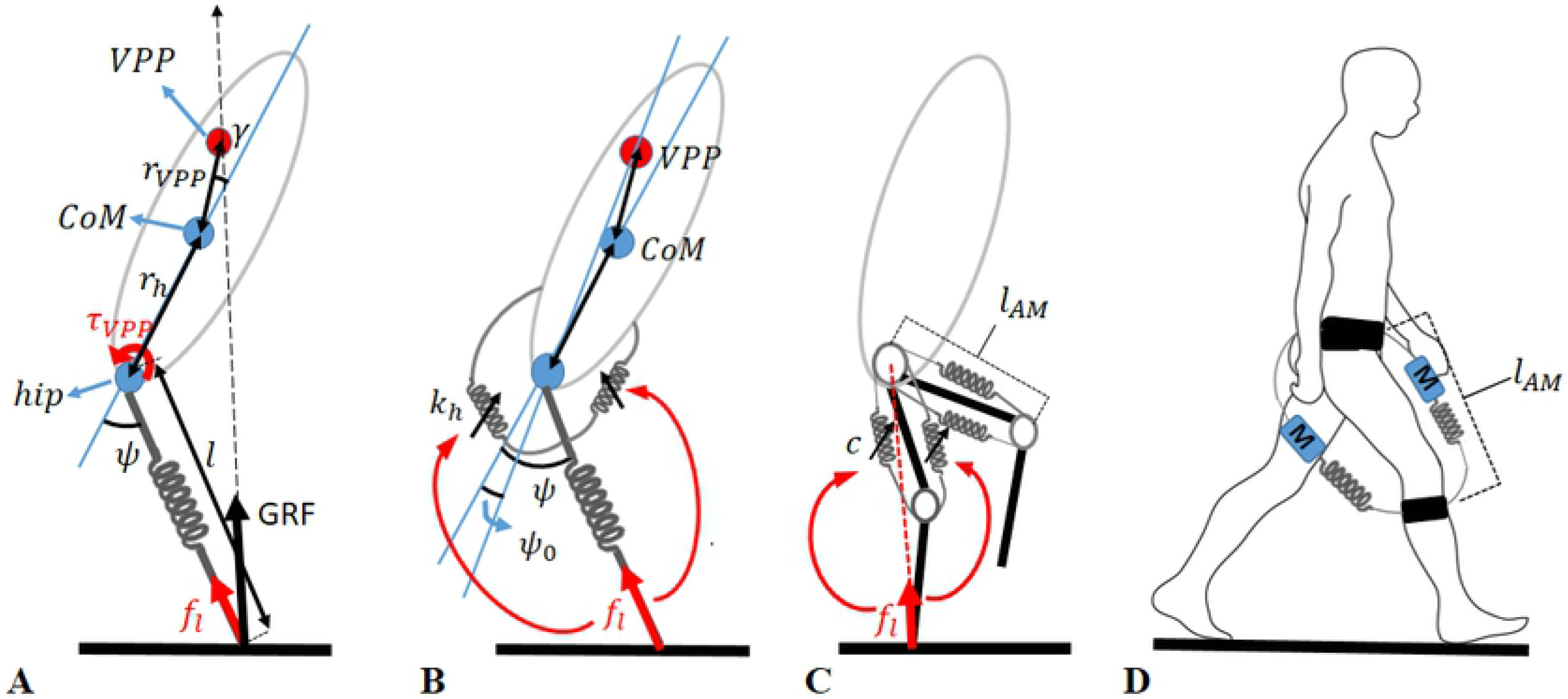
Evolution from template modelling to design of the BAExo for gait assistance. **(A)** The VPP (virtual pivot point) concept, shown in the TSLIP model wherein the leg force is always going through the VPP as a point on the upperbody. **(B)** The FMCH model as the mechanical implementation of the VPP concept, in which the leg forces are used as feedback signals to adjust hip spring stiffness. **(C)** The ABC (adaptable biarticular compliance) control method using the FMC model in the stance phase and passive biarticular thigh springs in the swing phase. **(D)** The ABC-based exosuit design schematic drawing, a basis for the BAExo.

##### The FMCH model

In [36] a new template model was developed to generate VPP using a variable stiffness which is called FMCH (force modulated compliant hip) model. In this model, leg force feedback was used to adjust hip spring in the TSLIP model. It was shown that for the range of joint angles observed in human walking, the FMCH can precisely approximate the predicted hip torques from the VPP model (Eq. 1).

Furthermore, this model could nicely predict the human hip torque *τ*_*h*_ in walking at different speeds [60] with the following simplified equation.

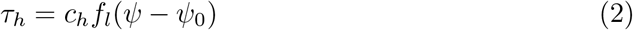

Here, *c*_*h*_ and *ψ*_0_ are the normalized stiffness (to body weight and leg length) and the rest angle of the adjustable hip spring, respectively (Fig. 1B).

##### The FMC model, an extension for segmented leg

Due to the segmentation of leg in humans, for using the GRF-based control (e.g., to control the exosuit) the FMCH model was extended to a biarticular level [51]. Since we use the GRF for controlling biarticular muscles and not the hip spring, we will use *FMC* standing for force modulated compliance and skip the “H(hip)” in FMCH. For this, a virtual leg was defined by a line from the hip to the ankle and the virtual hip torque was defined between the virtual leg and the upper body. To control this virtual hip torque both hip and knee joints should be controlled in coordination. In [43] it has been shown that with appropriate design of the thigh biarticular actuators in a bioinspired bipedal robot (BioBiped3), GRF direction can be controlled with minimum interference to GRF magnitude. For as much as the VPP concept is based on GRF direction control, the thigh biarticular muscles (simplified as adaptable springs in Fig. 1C) can be used to mimic human-like balance control in the segmented leg. Thus, the force of biarticular (artificial) muscle, given by this so called FMC controller, will be

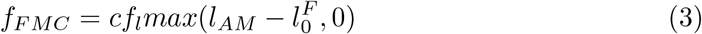

in which *f*_*FMC*_ is the force applied by the biarticular actuators (adaptable springs for RF or HAM artificial muscles), *f*_*l*_, is the axial leg force, and *c*, *l*_*AM*_ and 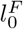 are the normalized stiffness, length and rest length of the artificial muscle, respectively. For the two biarticular thigh muscles, *c*, 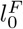 are the tunable control parameters which will be determined by the optimization method described in the following section. The *max* function represents the (muscle-like) unidirectional force generation of the actuators. In [51], we introduced this method to be applied for design and control of an exosuit with one actuator mimicking just HAM muscle (hip extensor, knee flexor). Based on the same argumentation of [43] the hip to knee lever arm ratio was set to 2 which generates the same effect as the FMCH with springy leg [51] and equalizes models of Fig. 1A and B. In this study we extend the method by 1) adding the second actuator (for RF), 2) adding a compatible swing leg control, 3) introducing an optimization based design method using OpenSim software and 4) implementing on the hardware setup. These extensions are described in detail in the following.

#### Swing phase control

Dean and Kuo with a simulation study demonstrated that coupling the hip and knee joints with biarticular springs can yield efficient swing motion with an appropriate ground clearance similar to humans [61]. They also showed that the knee-to-hip moment arm ratio has a significant effects on stability of gaits at different speeds. More recently, Sharbafi and his colleagues could predict kinematic and kinetic behavior of swing leg and muscle forces in human walking using a template model consisting of a double pendulum with combinations of biarticular (thigh) springs [52]. The outcomes of these studies are in line with former biomechanical studies, which emphasize the important role of the thigh biarticular muscles in the swing phase of human walking [62]. It was shown that the RF and HAM muscles contribute in the first and second halves of the swing phase, respectively [62]. These evidence support the ability to provide human-like swing leg motion with passive biarticular springs which also fits to our FMC control in stance phase (see Fig. 1C). Therefore, we used fixed springs (with constant *k*) for assisting hip and knee joints in the swing phase.

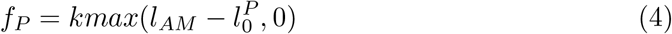

In this equation, *f*_*P*_, *k*, *l*_*AM*_, 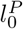 are the force applied by artificial muscle, stiffness, the length and the rest length of the artificial muscle for the passive mode, respectively.

#### ABC Control of BAExo in Walking

The ABC control is composed of adjustable biarticular springs with force modulated compliance for a time interval (*T*_*f*_) and fixed springs (with fixed stiffness *k*) for another time interval (*T*_*p*_). The following equation presents the formulation of ABC control for each leg.

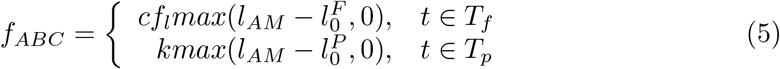

Here, *l*_*AM*_ is the length of the artificial muscle (AM) including the spring length and motor displacement. The reset length of AM is set differently for 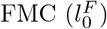 and passive mode 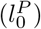.

For the HAM-artificial muscle, the time interval *T*_*p*_ starts with the takeoff (beginning of the swing phase, shown by *t*_*TO*_) and ends when the force generated by passive spring equals the force calculated by FMC. This moment of <u>f</u>orce <u>e</u>quality is denoted by *t*_*FE*_ and can be found when the following equality condition is held.

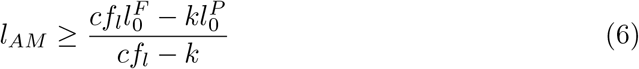

Then, *T*_*f*_ starts from *t*_*FE*_ to the next takeoff of the same leg. This switching rule is considered to generate continuous changing between the two phases of the ABC control.

For RF-artificial muscle, *T*_*f*_ starts by touchdown (beginning of stance phase, denoted by *t*_*TD*_) and ends at a certain time *t*_*lock*_ by locking the motor before beginning of the swing phase. Consequently, *T*_*p*_ is defined from this moment to the next touchdown. Determination of *t*_*lock*_ is described in Sec.. Roughly speaking, contribution of HAM-artificial muscle starts by passive spring (when the motor is locked) in the late swing and continues by switching to FMC in the stance phase, while the RF-artificial muscle starts with FMC in the stance and continues by switching to passive spring (by locking the motor at *t*_*lock*_) before swing phase starts. Our investigations showed that setting *t*_*lock*_ to a moment that the second peak of GRF appears (*t*_*P*2_) will result in more efficient and effective control (see description in Sec.). The influences of the spring stiffness and motors locking time on the serial spring rest length and the effective force are discussed in detail in the results section. The control architecture (including switching time and feedback signals are illustrated in the block diagram of Fig. 2. In this block diagram, the condition for start of passive mode and FMC mode has been shown by the (grey) dashed and solid lines, respectively. In the FMC mode, the reference force is calculated using GRF, AM-length and tunable parameters 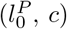, then SEA follows this reference force.

**Fig 2.**
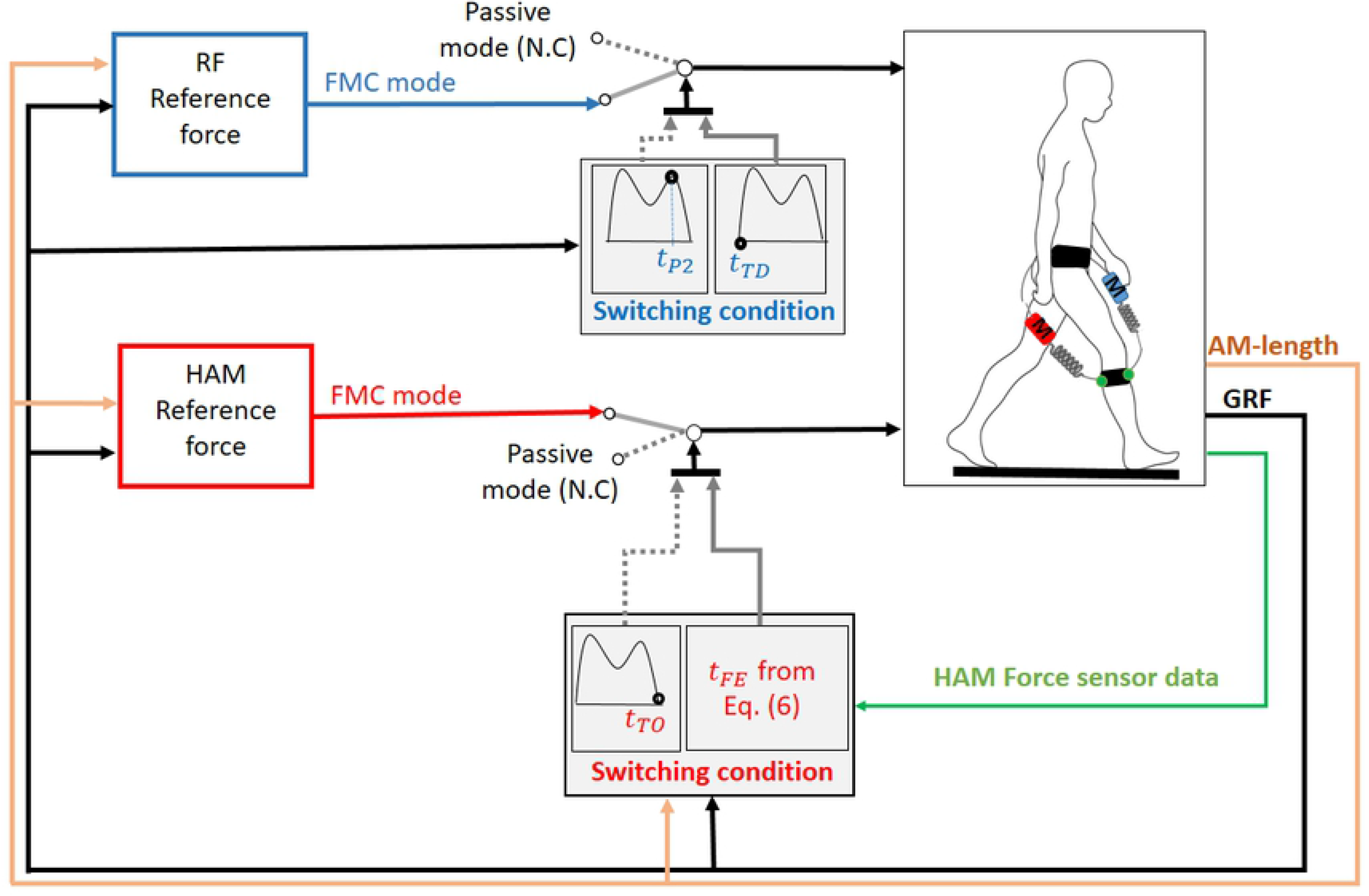
The ABC control block diagram illustrating the reference force generation and switching conditions. In the FMC mode, SEA follows reference force developed by modulating the artificial muscle stiffness with GRF feedback signal. In the passive mode, the motor is locked and the SEA acts as a passive spring. In this figure *t*_*P*2_, *t*_*TD*_, *t*_*TO*_ and *t*_*FE*_ are the moments that the second peak of GRF, touch down, toe-off and force equality (from Eq.(6)) occurs, respectively. N.C (stands for not connected) means in the passive mode, there is no reference force to be followed by the actuator.

Inspired by the positive force feedback of muscles [63], we consider about 40 ms delay (*t*_*d*_ = 0.040*s*) in the GRF signals. Interestingly, this biologically motivated delay results in a better match of FMC force with optimal force. In fact, this delay exists in the exosuit implementation duo to ground reaction force measurement and the settling time in the low level force control. All in all, timing active control (*T*_*f*_) is given by the following equations for HAM- and RF-artificial muscles

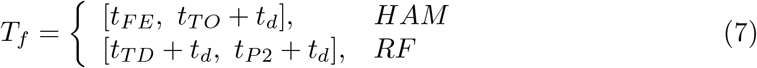

The passive time *T*_*p*_ in which the motor is locked will be the rest of the gait cycle. Eqs. (5) to (7) clearly show that the ABC control approach is independent of time, and the GRF and AM forces are used to detect the switching time and required forces.

### OpenSim model development for testing BAExo

For evaluating the applicability and performance of the proposed BAExo design using ABC control, we employed OpenSim software (https://opensim.stanford.edu). Due to difficulties in (human) experiment based validation of assistive devices’ design and control, the method of using neuromuscular simulation models is becoming more popular [5, 30, 64–66]. By reviewing several studies with this approach Grabke et al., stated that the musculoskeletal modeling-based approaches can be considered as alternatives to intuition-based assistive device design [67].

OpenSim is an open source software for developing experiment-based models of musculoskeletal structures and creating dynamic simulations of movement [68]. Using an open data-set in OpenSim, we added the BAExo to the human walking model and searched for the optimal actuation to minimize the approximated metabolic cost. Such a data-driven optimal force pattern is not generated based on a control principle, but it can be considered as a reference for designing the controller. For this, we applied the ABC control method to deliver forces close to the optimal ones. In the following, we first describe human walking data used in the OpenSim model and then discuss the optimization process and the ABC implementation on the BAExo.

#### Experiment-based OpenSim simulation model

The locomotion task in this study is walking at preferred speed (1.45 ± 0.15 m/s) for 6 healthy subjects (age 25 ± 5 years, height 1.86 ± 0.04, details in Tab. 2). This data-set is borrowed from [5] which is available at https://simtk.org/home/assistloadwalk. For each subject, we used 3 overground trials in which the kinematic data, ground reaction forces (GRF) and moments were measured. Using 8-camera optical motion capture system, the trajectories of 41 markers were collected at 100 Hz (Motion Analysis Corp., Santa Rosa, CA, USA). Also, the ground reaction forces and moments were collected at 2000 Hz from 3 floor-mounted force plates (Bertec Corp., Columbus, OH, USA). Because of differences in the preferred speed and stride length of subjects, in some trials, the data on one of the force plates was lost. Hence, the force data was not sufficient for modeling a complete stride in some trials. As a result, we analyze the experimental data for two single support and one double support phases. Considering the contribution of different (right and left) legs, this time domain is adequate for analysing the assistance from the exosuit and the corresponding muscles’ activities.

**Table 2.** Subject-specific experimental data from dembia2017simulating

| Subject | mass [kg] | height [m] | age |
| --- | --- | --- | --- |
| 1 | 112.4317 | 1.9177 | 27 |
| 2 | 89.2338 | 1.8796 | 20 |
| 3 | 86.9085 | 1.905 | 19 |
| 4 | 64.0353 | 1.8034 | 21 |
| 5 | 85.0668 | 1.828 | 31 |
| 6 | 67.1419 | 1.8288 | 32 |

We used OpenSim software (version 3.3) for simulations [68, 69]. A three-dimensional musculoskeletal model with 39 degrees of freedom and 80 Hill-type muscles has been used for our simulations. Eight degrees of freedom (bilateral ankle eversion, toe flexion, wrist flexion, and wrist deviation) which are not necessary for our analyses has been locked. The model was based on 21 cadavers and 24 young healthy humans [70]. Our simulation workflow was based on [5]. At first, the musculoskeletal model was scaled to match the anthropometry of each subject. Then, joint angle trajectories were calculated using OpenSim’s inverse kinematics (IK) tool. After that, OpenSim’s residual reduction algorithm (RRA) tool was used to reduce the residual forces (applied to the pelvis) resulting from small discrepancy between force plate data, marker data, and the musculoskeletal model. Finally, by using OpenSim’s computed muscle control (CMC) tool, muscle driven simulations of each trial have been generated.

#### Optimal control of BAExo

We utilized the CMC tool for designing and predicting the effects of ideal assistive devices. The two biarticular (HAM- and RF-) actuators, shown in Fig. 1, were implemented in OpenSim using *Path Actuator* (Fig. 3A). This device is defined to be unidirectional (developing just pulling force) and we added it in bilateral configuration (to both legs). The devices are implemented in OpenSim such that the exosuit only applies force in the sagittal plane. This is how the developed BAExo hardware setup works as well. For simplicity the introduced exosuit was massless with ideally perfect force tracking without power or force limitations.

**Fig 3.**
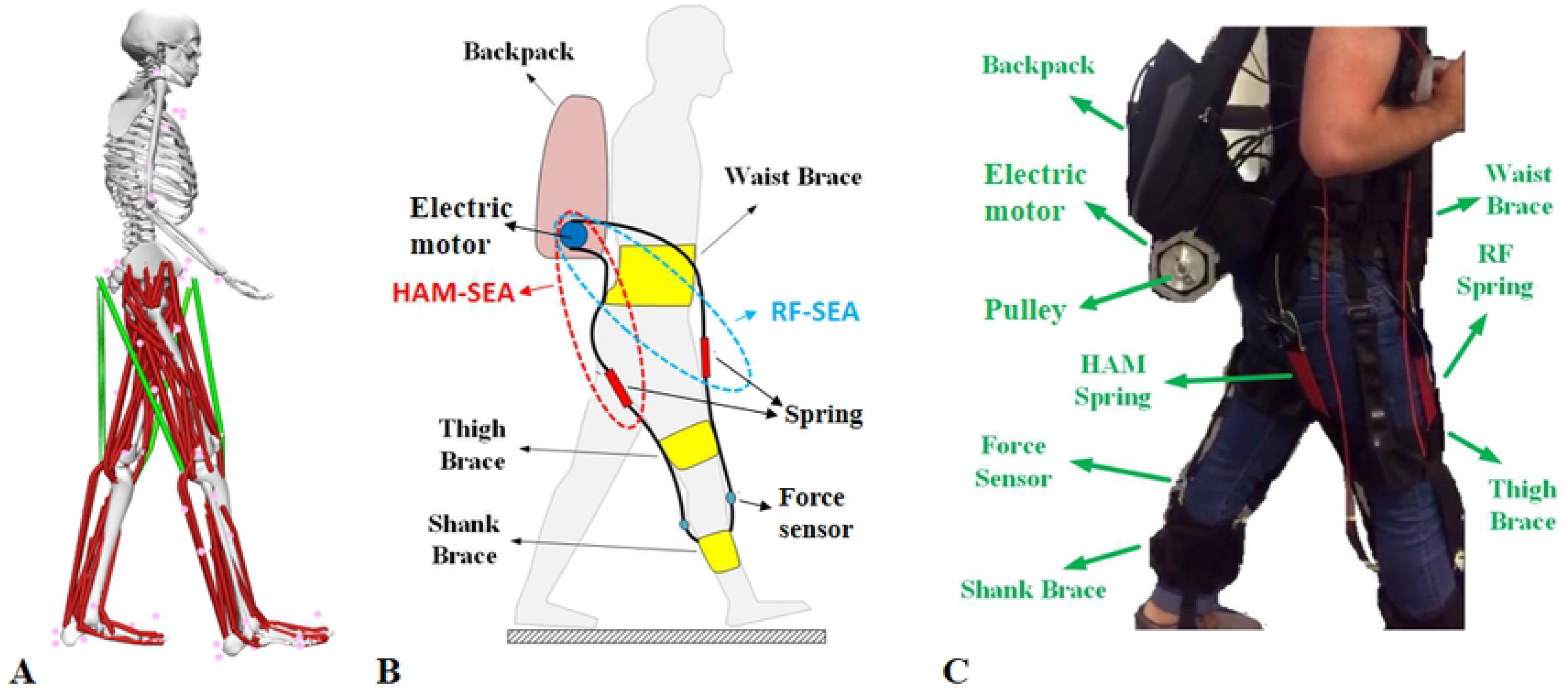
Implementation of the BAExo. **(A)** Implementation of the BAExo (exosuit with biarticular thigh actuators) in OpenSim. Red lines show muscles in the model and green lines indicate actuators in the model. **(B)** Schematic of BAExo design. **(C)** Picture of developed BAExo using SEA for both biarticular thigh actuators mimicking HAM and RF muscles in human body.

In order to minimize the muscle energy consumption at each time instant, we can use the instantaneous muscle activation signals. As the generated motion (and consequently muscle length changing rate) is fixed and determined by the human experimental data, muscle activation could be an appropriate signal to approximate and minimize the consumed energy. Therefore, the CMC’s cost function, *J* at time instance *t* is defined based on two terms: body effort, and simulation (modeling and measurement) error. This cost function (*J*) is borrowed from [5]. Considering *M* as the set of lower limb muscles, the effort term is calculated by summation of square of individual muscle instantaneous activation *a*_*i*_ (*t*) for *i* ∈ *M*. This OpenSim model considers a set *R* of so called *reserve* and *residual* actuators to generate forces/moments *f*_*j*_(*t*) for *j* ∈ *R* which compensate for measurement and modeling errors (see details in [5]). *Reserve actuators* added small torque about each joint to compensate for unmodeled passive structures (e.g., ligaments) and potential muscle weakness. *Residual actuators* apply the residual forces to minimize the effects of modeling and marker data processing errors. The utilized cost function at each time instance *t* is given by

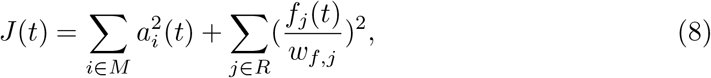

in which *w*_*f,j*_ denotes the constants for weighting the compensating actuator forces [5]. In the assisted mode, the CMC’s cost function *J*_*A*_ includes the force *f*_*k*_(*t*) applied by the exosuit actuators in addition to the previous terms. Defining *E* as the exosuit actuator set and *k* ∈ *E* the updated cost function will be

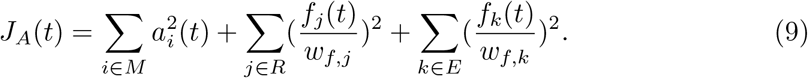

Here, *w*_*f,k*_ weight is set to a large value (1000 N) for lowering the penalty of using actuators instead of muscles. We applied the same kinematic and kinetic (GRF) dataset (explained in Sec.) to both assisted (with BAExo) and unassisted models. Thus, the net joint moment in assisted and unassisted modes is conserved and the provided supportive effort in the exosuit could potentially yield lower muscle instantaneous activation. This could result in metabolic cost reduction. We employed the cost function *J*_*A*_ to find the optimal force at each time instant. The resulted control patterns will provide maximum reduction in metabolic cost with the BAExo, while maintaining the kinetic and kinematic behaviour. This means that theoretically, applying these patterns to the BAExo will help the individual subjects to consume the minimum metabolic energy.

#### ABC control in OpenSim

Following predefined patterns obtained by optimization in Sec. does not have any fundamental difference to the trajectory-based methods (see Sec.). Instead, here we try to use this optimal signal as a reference to tune our bioinspired ABC controller. This way, a closed loop control with GRF as a feedback signal can be applied to real exosuits which in principle could outperform the robustness and adaptability (e.g, perturbation recovery or adapting to changed gait conditions) of the feedforward (trajectory-based) control.

In Sec. we described the ABC control method which can prescribe the appropriate force in biarticular thigh artificial muscles in both stance and swing phases. The stiffness and rest length of the fixed and adaptable springs in Eq. (5) will be identified by fitting the ABC forces to the optimal reference trajectories using MATLAB curve fitting toolbox. Using the optimal patterns, we also found that the second peak of the GRF is a perfect moment to be selected as the locking time *t*_*lock*_ for the RF-artificial muscles. This way, the dependency to time is removed by using the GRF which was already measured and used in the FMC control (see Sec. for details).

For implementing the ABC controller in OpenSim, first, we calculated force profiles with Eq. (5) using kinematic and ground reaction force of the unassisted trial. Then, using OpenSim’s CMC tool this force profile is applied to the model and the resulting metabolic cost is calculated. The main constraining assumption of these simulation studies is that subjects walked with the same kinematics and ground reaction forces in both assisted and unassisted conditions. Studies on soft exosuit reported relatively small changes in kinematics and kinetics with assistance [71]. Hence, assuming fixed kinematic and kinetic behaviour is acceptable.

### Design and control of BAExo hardware setup

In addition to developing OpenSim simulation models, we also designed and manufactured the hardware setup of the BAEXo. We directed a pilot experiment with one 27 years old male subject with 72kg weight and 178cm height. The experiment goal was to verify the functionality of the designed exosuit and the proposed ABC control approach. The subject was walking on an instrumented treadmill at 1.1 *m/s* as his preferred speed.

The exosuit is composed of a wearable part and an actuation system. The textile components of BAExo consist of a waist brace, two thigh braces and two shank braces (see Fig. 3). There are also HAM and RF serial compliance that consists of a rubber band, Bowden cables and force sensors (see Fig. 3C). The portable actuation system with 2 electric motors will generate assistance force during walking. The actuation components (motors, gears, motor drivers, pulleys and wires) and electronic boards are placed in a backpack (see Fig. 3C). Detailed specification of the BAExo design is listed in Table. 3.

**Table 3.** BAExo design specifications.

| part | mass[kg] | description |
| --- | --- | --- |
| Waist brace | 0.55 | Textile |
| Thigh brace | 0.23 | Textile |
| Shank brace | 0.17 | Textile |
| HAM SEA | 0.35 | stiffness, 3.3 kN/m |
| RF SEA | 0.31 | stiffness, 1.1 kN/m |
| Backpack | 1.7 | driver , box , wire |
| Motor | 0.48 | Maxon, Graphite Brushes, 150 Watt |
| Gear | 0.46 | Maxon, Planetary Ceramic Gearhead |
| Pulley | 0.086 | diameter,50mm |
| Force sensor | 0.01 | KM10Z, 200N |

For reducing the weight of the exosuit, we used one motor (with augmented gearbox) for each leg, as shown in Fig. 3B,C. These motors are used to pull two serial elastic rubber bands. Therefore, using these SEA arrangements, the same motor is employed for generating the required force in both biarticular thigh artificial muscles. Therefore, by rotating the electric motor in different directions, HAM- or RF- artificial muscles produce forces. Bowden cable is used to transmit the force from the motor to biarticular elastic elements (HAM, RF; Fig. 3B and C). On the top side, the Bowden cable sheath is connected to the frame of the pulley cover and the inner cable is attached to the pulley. The other end of the Bowden cable sheath is connected to the fixed point on the bottom of the waist brace and the inner cable is connected to the rubber band. The lower side of this serial elastic element is connected to the force sensor that is fixed to the shank brace. A thigh brace is used to align the serial elastic element with human limbs. These components help to concentrate the SEA force in the sagittal plane and to keep the knee lever arms of both artificial muscles around 4 cm.

Two force sensors (KM10Z, 200N) are attached to the shank brace to measure HAM and RF SEA’s froce signals. The vertical GRF was measured individually for each limb using the instrumented treadmill (ADAL-WR, HEF Tecmachine, Andrezieux Boutheon, France) as a feedback signal for the FMC approach. Different (stance and swing) phases of the gait was detected based on the GRF data. Combined with the actuator position signals, measured by the encoders mounted on the back of DC motors, the hybrid ABC control was implemented in the hardware setup. In addition to supporting actuation and sensing, the system provides real-time feedback control and data logging at a frequency of 1 kHz.

## Results

This section describes the simulation and experimental results of gait assistance with our bioinspired designed and controlled exosuit. First, the design of the ABC control (matching the optimal trajectories) of the BAExo in the experiment-based OpenSim model is described and demonstrated. Then, the quality of this control is evaluated based on the metabolic cost reduction (with respect to unassisted walking) and exosuit power consumption and the comparison to the optimal solution. Since the experimental data (used in OpenSim models) is collected using three force plates, part of the GRF data is missing for one of the stance phases, due to miss-placement of the foot for some trials. To have consistent calculations for all subjects, we removed the incomplete force data and demonstrated the remaining results which included two single supports and one double support (from about 20% to 100% of the gait cycle). Finally, in order to demonstrate the applicability of the proposed method, the outcomes of a pilot assisted walking experiment (with the BAExo) on the instrumented treadmill are presented.

### Optimization based ABC control in OpenSim

In order to find the control parameters of the ABC approach, first, we need to identify the optimal force patterns which are calculated using the method introduced in [5] (see Sec. details). In this section, we separate the design process of the HAM- and RF-artificial muscles. In Fig. 4, the optimal force for the HAM-artificial muscles is shown by gray colour. In the late swing phase, the optimal HAM-artificial muscles starts to contribute. This contribution can be provided by a passive biarticular spring. The optimal stiffness in the simulation is selected according to optimization force in the *T*_*P*_ time interval. We found that 15 – 20 *kN/m* is an appropriate stiffness range for HAM-artificial muscle approximate the optimal solution for different subjects. In the stance phase, the second peak of the optimal force can be perfectly approximated by the FMC (the blue curve). This result is consistent for all subjects. Combination of passive biarticular spring in the swing phase and the FMC in the ABC framework (the red dashed curve) nicely predicts the optimal pattern. This result is valid for most of the subjects. In the proposed ABC control for HAM-artificial muscles, the motor is locked during swing phase (interval *T*_*p*_) and the AM behaves like a passive spring in late swing as shown in Fig. 4B. From beginning of stance phase, the FMC predicts increasing force which is opposite to the generated force by the passive spring. As soon as these two forces are intersecting, the ABC switches to FMC which means stretching the spring by moving the motor (see Fig. 4B). Interestingly, the required movement to follow the FMC is small and slow which could result in low power requirement.

**Fig 4.**
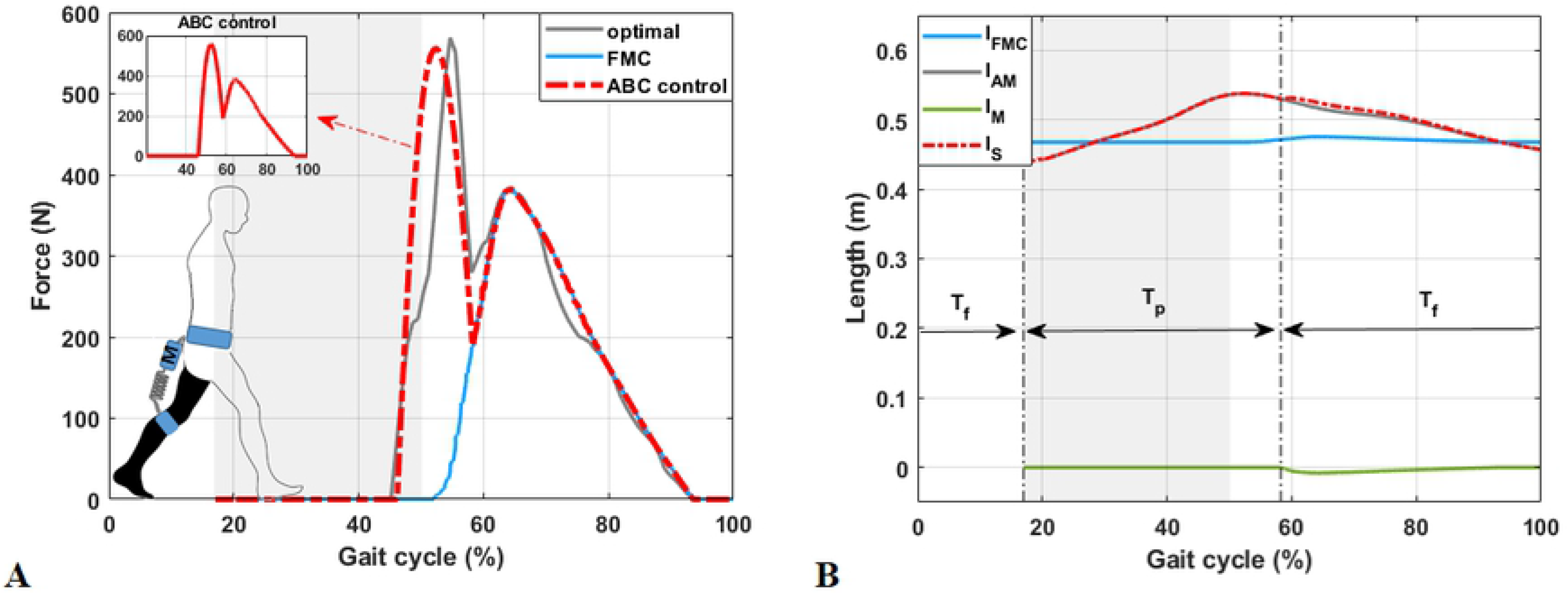
HAM-artificial muscle: force and length. **(A)** The HAM-artificial muscle optimal force pattern for the left leg and its approximation by the ABC controller (using SEA) in the OpenSim walking model. **(B)** Displacement of the motor (*l*_*M*_) and the length of the serial spring (*l*_*S*_) and HAM-artificial muscle (*l*_*AM*_). Here, the *l*_*FMC*_ is the prescribed length of the serial spring to apply FMC force, which is not followed after *t*_*lock*_. *T*_*p*_ and *T*_*f*_ are respectively, passive mode and FMC operation time intervals, used in ABC control method. The shaded region shows the swing phase of the gait cycle. The gait cycle starts with the touch down of the right leg.

For the RF-artificial muscles, we used an SEA for force generation. Similar to human RF muscles, here the optimal force starts at mid-stance (about 35% of the the gait cycle, as shown in Fig. 5). By tuning the appropriate rest length and normalized stiffness, the FMC can appropriately predict the optimal force in the stance phase including the first peak (see Fig. 5A). The second peak in the optimal pattern is not consistent with the biological evidence of humans RF muscle actuation in walking [62]. This means that the RF-artificial muscles can reduce the activation of other muscles. We tried to find the best fit for the optimal RF-artificial muscles force with minimal effort. The blue curve in Fig. 5A, shows that with only FMC the artificial muscle force will be limited to the stance phase. The green curve shows the best fitted passive design (just spring, without motor) to the optimal solution which cannot contribute to the second half of swing phase. In ABC control architecture, locking the motor (equal to switching off the motor for non-backderivable motors), turns the FMC into a passive spring which can still contribute in the swing phase. This way, part of the optimal force can be provided without energy consumption (investigating the locking time *t*_*lock*_ is described in the following).

**Fig 5.**
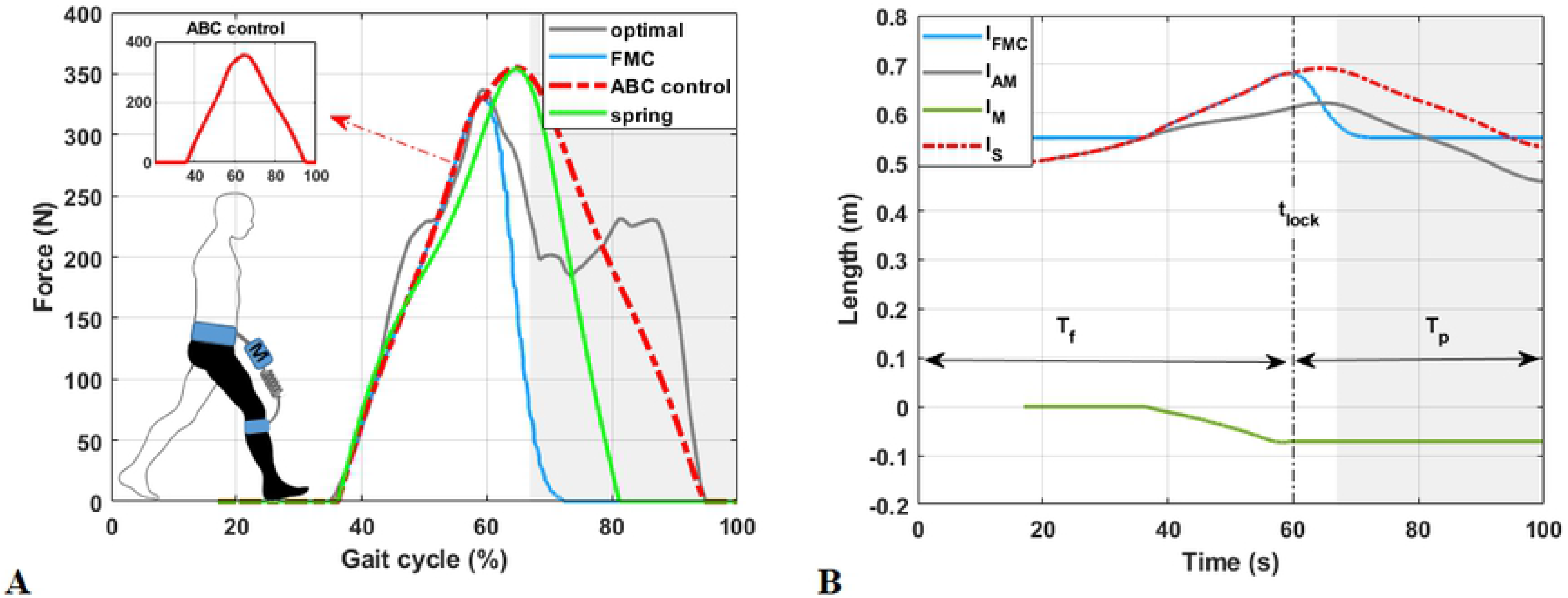
RF-artificial muscle: force and length. **(A)** The RF-artificial muscles optimal force pattern for the right leg and its approximation by the ABC controller (using SEA) in the OpenSim walking model. **(B)** Displacement of the motor (*l*_*M*_) and the length of the serial spring (*l*_*S*_) and RF-artificial muscle (*l*_*AM*_). Here, the *l*_*FMC*_ is the prescribed length of the serial spring to apply FMC force, which is not followed after *t*_*lock*_. *T*_*p*_ and *T*_*f*_ are respectively, passive mode and FMC operation time intervals used in ABC control method. The shaded region shows the swing phase of the gait cycle. The gait cycle starts with the touch down of the right leg.

The movement of the motor and the spring for RF actuator in the prescribed ABC control (dashed red curve in Fig. 5A) is depicted in Fig. 5B. The motor only moves from mid-stance to *t*_*lock*_ resulting in an increase in the spring stored energy. This enhances forward movement of the upper body and swing leg in the late stance and swing phases, respectively. The blue line in this graph illustrates the spring length if the FMC was implementing completely (without locking in between). The difference between the spring (*l*_*S*_) and the RF-artificial muscles (*l*_*AM*_) lengths is constant after *t*_*lock*_.

In order to better understand the ability of the proposed ABC control within the SEA arrangement, we investigated the effect of the serial spring stiffness and motor’s locking time *t*_*lock*_ in Fig. 6. The *t*_*lock*_ influences the contribution of the motor (consumed energy) and the rest length of the spring which also affects the duration and the magnitude of the RF-artificial muscles contribution in the swing phase. If we lock the motor shortly after it generated the peak force (the GRF second peak) (shown in Fig. 6A), tracking optimal pattern with the ABC controller is improved. However, the motor needs more energy and power as it works longer and needs to stop and return in the opposite direction, immediately. Locking before reaching the peak force results in decreasing motor energy and power, while lowering the quality of tracking the optimal pattern by diminishing the first peak. Our simulations show that locking the motor when the second peak of GRF occurs, is a perfect compromise between human metabolic rate reduction and motor power consumption. Such timing does not require a sudden change in motor movement direction and yields in providing part of the RF-actuator contribution in the swing phase. This also supports the time-independent control described in Sec..

**Fig 6.**
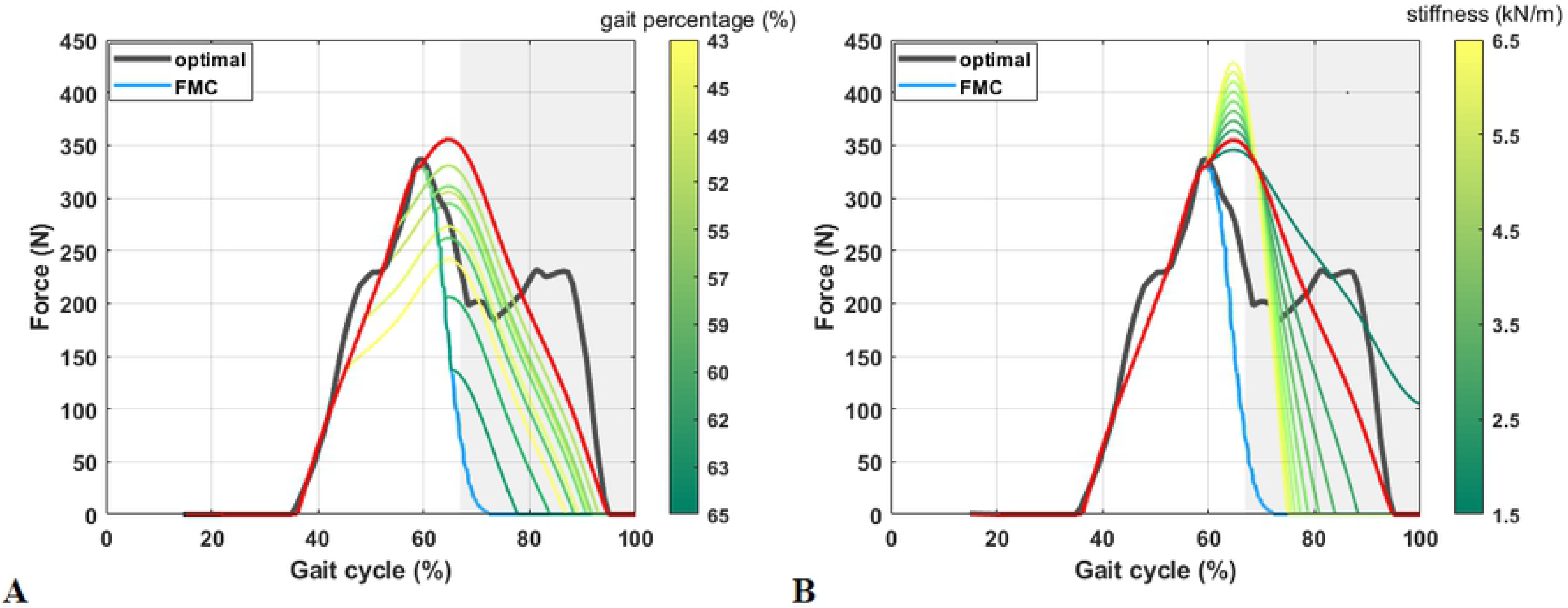
Parameter tuning in RF-artificial muscles for control of the BAExo. **(A)** The effects of the motor’s locking time *t*_*lock*_ on the ABC force patterns. Here, the colour bar shows the percentage of the gait cycle for *t*_*lock*_ and the stiffness is equal to 2500N/m. **(B)** The effects of the SEA spring stiffness on the ABC force patterns. The colour bar shows the stiffness in *kN/m* and the occurrence of the GRF second peak is selected to set the *t*_*lock*_.

Increasing stiffness of the serial spring, enlarges the magnitude of the RF-artificial muscles force (consequently the peak value), as shown in Fig. 6B. With larger stiffness, we need less displacement of the motor to create the same force with the FMC which means less lengthening in the serial spring. This results in shorter contribution of the passive spring in the swing phase. Therefore, by decreasing stiffness, the stored energy and lengthening of the spring (as well as motor’s displacement) are increased which results in applying force for a longer period in the swing phase. Fig. 6B shows that 2500N/m is an appropriate spring stiffness which can approximate the optimal solution the best. In this figure, the moment of reaching the second GRF peak is selected as *t*_*lock*_.

### Metabolic cost reduction

We calculated the metabolic cost by integrating the instantaneous whole-body metabolic rate over one walking step (one single and one double support) using the *Umberger*2010*MuscleMetabolicsProbe* in OpenSim 3.3 [5, 72]. According to the subsection 2.2.1, due to the quality of our data, we focused on analyzing one double support and two single support phases. Thus, to estimate an average whole-body metabolic rate, we averaged the instantaneous whole-body rate over half a gait cycle (considering the approximate mediolateral symmetry of walking). To assess the effect of this approximation on our results, we compared the percentage of metabolic energy reduction of using complete gait cycle and half gait cycle in some trials in which complete gait cycle’s data are available. We found the negligible difference in our results between using a complete gait cycle and a half-gait cycle.

Similar to Optimal controller, the BAExo with the ABC control can also reduce required energy for walking at preferred speed. The metabolic cost reduction and the exosuit energy consumption of the optimization-based and the ABC control methods are shown in Fig. 7, 8. Comparison between percentage of the metabolic cost reduction for different subjects in Fig. 7, shows that our control method can effectively reduce the metabolic cost of walking by about 60%-80% of the optimal method. Although the reduction in the ABC control is less than the optimal approach (which was expected), it can provide a bioinspired white box controller, instead of black box method of giving a time-based trajectory, obtained by optimizing for a specific data set.

**Fig 7.**
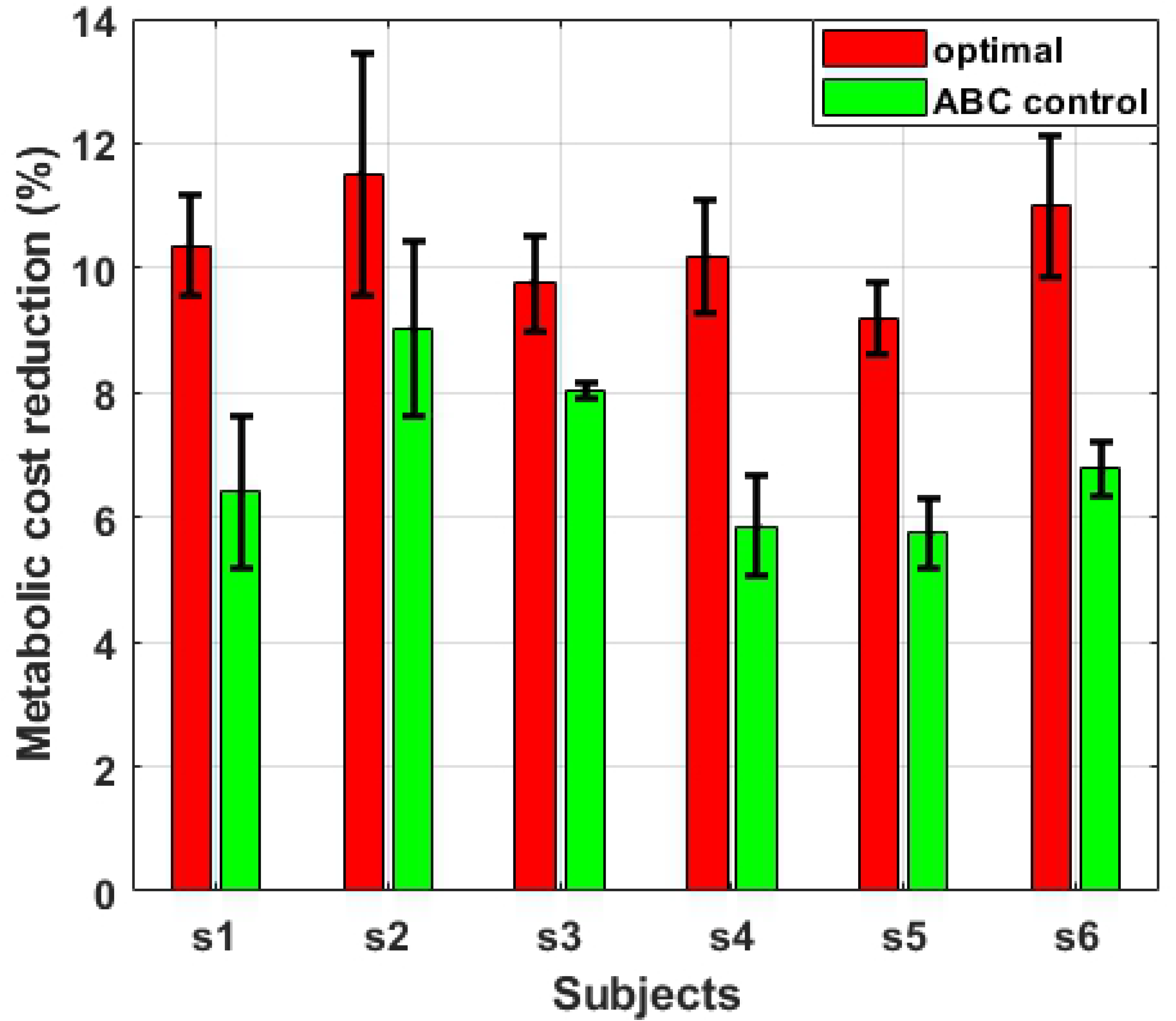
Simulation results: Percentage of metabolic cost reduction compared to unassisted mode. Percentage of metabolic cost reduction for the ABC and the optimal controllers for different subjects (*s*_1_ to *s*_6_), compared to unassisted mode. The mean and the standard deviation of three experimental trials for each subject are shown by the bar height and the error-bar, respectively.

**Fig 8.**
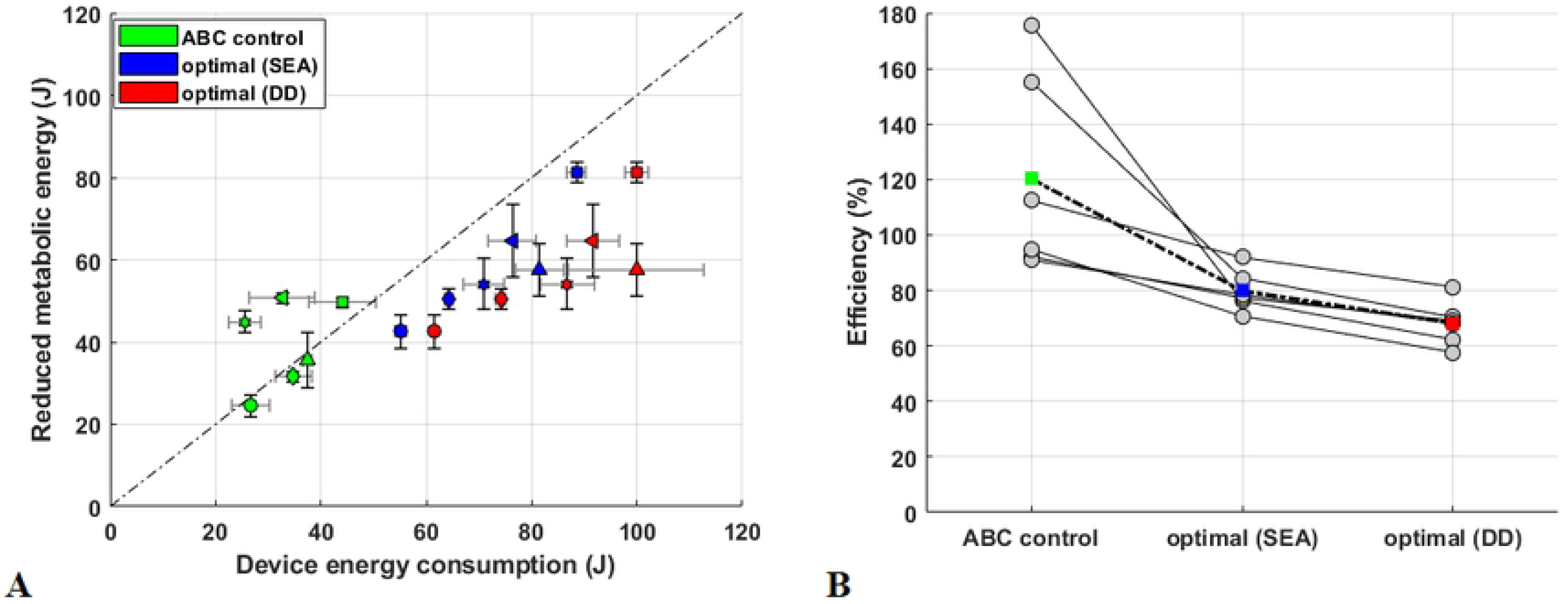
Simulation results: Metabolic energy reduction, device energy consumption and efficiency of different controller. **(A)** Metabolic energy reduction compared to unassisted mode and device energy consumption for the ABC and the optimal controllers for different subjects (showed by different marker). The mean and the standard deviation of three experimental trials for each subject are shown by the bar height and the error-bar, respectively. **(B)** The efficiency (defined by the ratio between metabolic cost reduction and consumed energy in the device) of the ABC and the optimal controllers for the BAExo in OpenSim. Note that, both optimal DD and optimal SEA are in the actuation level of applying the same optimized force patterns on the exosuit. Circle and square markers show the mean efficiency for each subject and the mean efficiency for each method respectively.

To provide a basis for comparing the efficiency of different methods, we illustrate the reduced metabolic energy (in Joule, not in percent) in the human subjects body versus the consumed energy in the exosuit in Fig. 8A. As the ABC control also benefits from compliant design of the actuators (SEA), we also used a compliant design for the optimal solution to reduce the consumed energy in the BAExo. Note that, both *optimal DD* and *optimal SEA* are in the actuation level of applying the same optimized force patterns on the exosuit which will not affect metabolic cost reduction, shown in Fig. 7. As demonstrated in Fig. 8A, the results of optimization with direct drive (DD) are all below the line with a slope of one, representing 100% efficiency. By adding the serial compliance, the consumed energy is reduced by 15%. The comparison between ABC control and the optimal solution with SEA, shows that the ABC is in average more efficient. To facilitate realizing this comparison, the efficiency (defined by the ratio of the saved energy in human body to consumed energy in the exosuit) is depicted in the separated graph (Fig. 8B). This graph shows that, not only the average efficiency of different subjects, but also control efficiency for each subject is lower for ABC compared to (SEA-equipped) optimal case. It is important to mention that the efficiency above 100% in Fig. 8B is not irrational as the BAExo design benefits from compliance and biarticular structures (see discussions in Sec. for more details).

### Exosuit control implementation

The ABC control approach -represented in block diagram Fig 2- is implemented in the BAExo hardware setup. The control sequence (timing), the GRF and artificial muscle forces are depicted in 9. In the simulations, two separate actuators were used to implement ABC control method in RF- and HAM artificial muscles (AM). The employed control sequence, is shown in Fig. 9A. This is a typical pattern, developed based on the double-hump GRF signal shown Fig. 9B, and might slightly change step to step. In other words, this is not a fixed time-based pattern. The proposed control approach will result in AM force profiles, illustrated in Fig. 9C. As can be seen in this graph, the two AMs are not actuated simultaneously except in a short period around 40% of the gait cycle. Note that, despite activating FMC control for the RF artificial muscle from beginning of stance phase, it will not generate force until mid-stance. Similarly, the HAM artificial muscle will be slack in the late stance. This means that both AMs can be controlled by one motor connected to the two springs (rubber bands), as explained in Sec.. This simpler (and lighter) arrangement could also result in simpler control protocol. By merging the RF- and HAM-artificial muscle force generation in Fig. 9A, the motor control pattern for the experimental setup (with one motor) will be given by the sequence shown in Fig. 9D. The electric motor will start to stretch the HAM artificial muscle as soon as the generated force (measured by the force sensor) is equal to the desired FMC-based desired force. This is the force equality moment which was explained by Eq. (6). After passing the mid-stance, the HAM force will converge to zero while the RF force will start to increase. Therefore, after mid-stance the motor will rotate in opposite direction to stretch RF artificial muscle and follow the FMC for this AM. This will continue until the second peak of GRF (shown in Fig. 9E) which occurs slightly before take-off moment. At this moment, the motor will be locked to switch to the passive mode. This simplified ABC control method is developed based on the GRF, depicted in Fig. 9E and will result in AM forces drawn in Fig. 9F. This figure demonstrates that the force-based control of the adaptable biarticular compliance (ABC) method could nicely match simulation results as well as human thigh artificial muscle patterns in walking [62].

**Fig 9.**
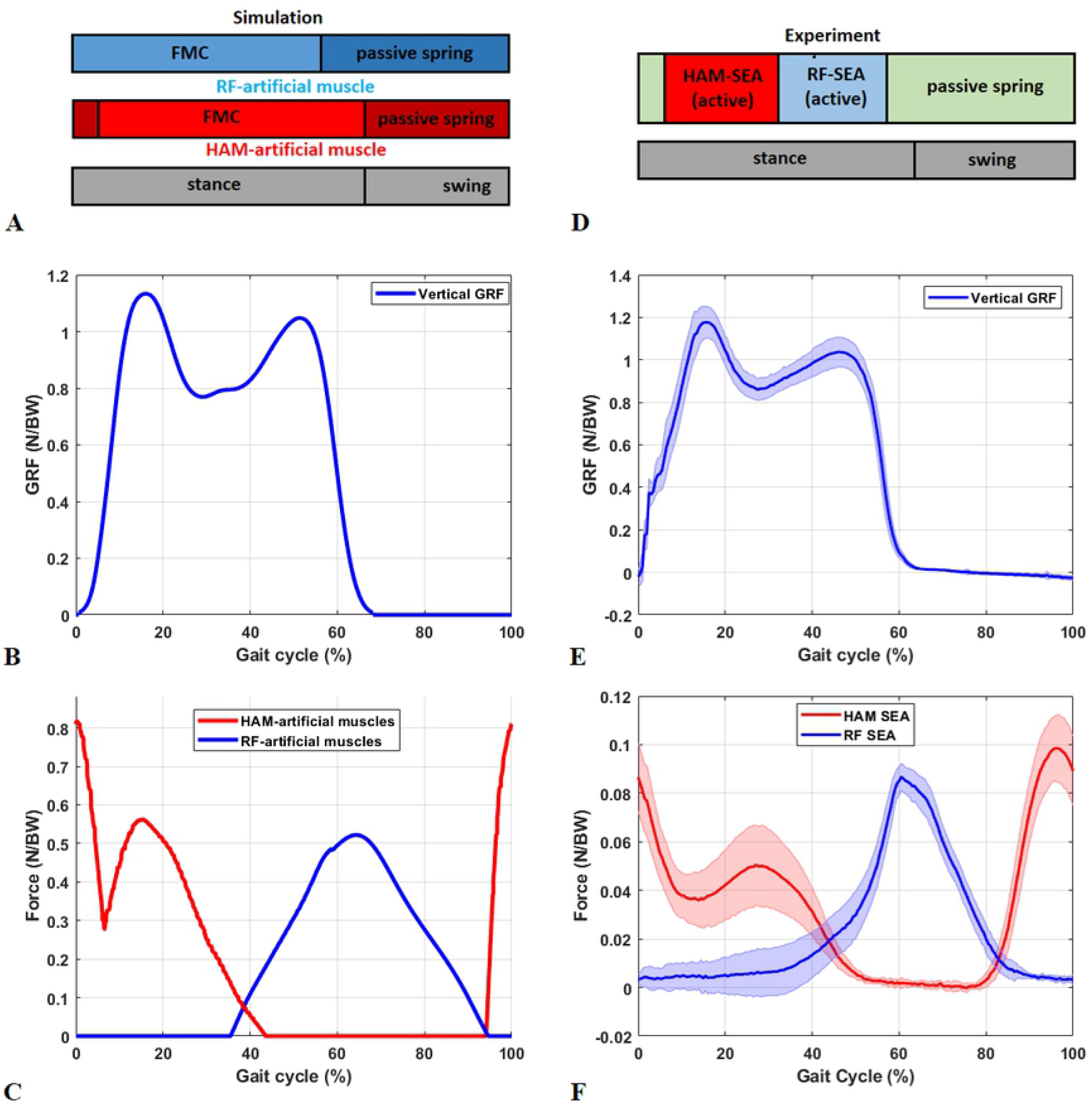
Comparison between simulation (A-C) and experimental (D-F) results: **(A,D)** The ABC control timing scenarios for one stride. **(B, E)** The ground reaction force patterns. **(C,F)** HAM-(Red) and RF (blue)-artificial muscles force profiles.

Roughly speaking, in the stance phase, the desired trajectories will be followed by a low level force control using the force sensors and in the swing phase the motors will be locked using position control. The attached video (S1 Video) shows the motor movement and the functionality of the BAExo assisting a subject in walking on an instrumented treadmill. Based on the subject self explanation, he feels assistance and a clear difference to the transparent (zero torque, applied by disconnecting the rubber bands from shank and switching off the motors) mode. The force patterns (GRF and the AM forces) look normal and as expected (Fig. 9). This experiment was to validate applicability of the control and design method for walking assistance. More quantitative (Oxygen consumption, EMG and kinematic) measurements with more subject are required to identify the assistance level.

## Discussion

In this study, we introduced BAExo as a new exosuit with innovative contributions in different levels of design and control: <u>1) Underlying control concept:</u> we developed a human-inspired control method instead of the common trajectory based control of the assistive devices [73]. <u>2) Actuation level:</u> implementing **compliant biarticular** actuation with SEA arrangement is proposed in BAExo design. Combination of biarticularity and compliance, as two bioinspired design features legged systems, are rarely applied to assistive devices [35]. <u>3) Feedback signal:</u> using the ground reaction force as a sensory feedback for control in locomotion which was inspired by biomechanical studies [41, 49] is applied for the first time in a soft wearable exosuit.

We investigated the proposed ABC design and control idea by human experiment-based simulations in OpenSim and preliminary experiments on human assisted walking with our recently developed BAExo robot. In order to facilitate identifying an optimal design and control for a new assistive device with respect to improving metabolic cost reduction in human locomotion, neuromuscular simulation models are advantageous. In simulation based analyses of gait assistance, we assume that the kinematic and kinetic behavior stays the same after wearing the exoskeleton as demonstrated in [71]. We accept the discrepancy between experimental and simulations especially in predicting humans’ reactions to external forces (e.g., from the exosuit). Nevertheless, such kind of neuromuscular experiment-based models (e.g., OpenSim models) are the best known software tools for proof of concept before experimental investigations. Furthermore, by using such simulation models, one can examine the effects of assistive devices on forces and metabolic consumption of individual muscles which is extremely difficult by experiments [5]. Hence, first we used the simulation model of gait assistance to evaluate our proposed approach (compared with optimal solution resulting in maximum reduction in metabolic cost). To validate the applicability of the method, we also implemented it on the hardware setup to show its functionality in interacting with humans. In the following, we discuss the outcomes of the simulations and experiments in light of the three above-mentioned contributions.

**1) underlying control concept:** Our FMCH-based control [36, 51] of the BAExo has a significant difference to the state-of-the-art methods of controlling assistive devices. Instead of replicating the outputs of the human locomotion control system e.g., joint torques, here we try to discover the underlying control principles for synthesizing locomotor functions. Recently, reflex-based control (introduced by Geyer and Herr [48]) was utilized for gait assistance [31, 66, 74]. Neuromuscular models [48] are useful biologically inspired tools for developing model-based control of assistive devices [30, 74, 75] which could potentially address the adaptability of the controller for different conditions. Nevertheless, the main barrier for using such models to control assistive devices is their complexity and a large number of parameters to be tuned [31, 74]. As the neuromuscular control in human locomotion is too complex and is not well understood [32], we employed a simplified model prescribed by the Template & Anchor concept [33]. Such kind of template-based control is recently employed for bioinspired legged locomotion control [76] and also assistive devices [19]. In that respect, we selected the FMCH model [36], which was developed to explain human balance control based on the VPP concept [50]. By use of biarticular thigh actuators with appropriate lever arms, an extension of the FMCH model for a segmented leg was presented [51]. This way, we translated a biomechanical template model of human balance control to a practical anchor level as a core concept for design and control of the so called BAExo. In the here here presented approach we combined this model with the passive biarticular spring model for human-like swing leg control (the DPS model in [52]). Therefore, the first and maybe the most important novelty of this study is introducing a new (template-based) approach for design and control of exosuits (for both stance and swing phase). If the fundamental control principles of human walking is correctly identified, we expect to have more harmonized interaction between the assistive device and human body. As a result, we expect to have improved robustness against perturbations and uncertainties which was not investigated in this study. Our preliminary experiments with BAExo partially supports robustness against walking speed changes. In a simple experiment we found adaptation of the force patterns when the treadmill speed was changing (not shown). Since such changes in gait characteristics will be reflected in the GRF patterns, our ABC control approach has sufficient sensory information to adapt. **2) Actuation level:** The force (GRF) modulated compliance model which is based on the VPP concept was already applied for control of LOPES-II exoskeleton in our previous studies [37, 38]. In addition to lack of swing phase control in the LOPES experiments, for implementing the FMC (GRF-based control) in such a rigid exoskeleton, the two single joint actuators were used to emulate the biarticular actuation. Therefore, part of the advantages of the proposed method were not met. Developing an exosuit with compliant biarticular actuators is our solution to regain these benefits. With biarticularity, transferring energy from one joint to another enables the system to reduce energy consumption [35]. Moreover, adding compliance in a serial elastic actuation mechanism provides further advantages such as, recoiling energy, increasing robustness (e.g., at impacts), reducing peak power and required torques of the electric motors. OpenSim simulations demonstrated the ability of the proposed mechanism to reduce metabolic cost up-to 12% with the optimized solutions. In these simulations, we assumed that the kinematic and kinetic behavior of human subject did not change. Definitely, adaptation of human subjects to the exosuit’s force could improve the control quality and may result in higher metabolic cost reduction. Experimental results show that the generated force patterns in the biarticular thigh actuators are similar to their biological counterparts (RF and HAM muscles). This supports the control concept, and the effects on the metabolic cost in real experiments could be tested in the future.

In our experiments, we tested 3 different stiffness values and we found that the subject feels more comfortable with the lowest value for the RF-artificial muscles and the highest value for HAM-artificial muscles. Since the optimal design and control parameters (in Eq. (5)) are subject-dependent, we do not expect to find similar values for the optimal solution in simulations and the experiments. Another important difference emerges from imprecision in the position and compliance of the attachment points (due to the soft tissue and movement between braces and the body which are not modeled). For example, different lever arms yield finding dissimilar spring stiffness values to generate the same joint torques, although we kept similar hip to knee lever arm ratio (close to 2) in both cases. Therefore, the simulation study can support the validity of the proposed method and the calculated parameters (e.g., spring stiffness) can be used as initial guess for the experimental study. Nevertheless, the optimal design and control parameters must be found separately in experiments with individual human subjects. Moderating optimal values, found from modeling analyses for the developed assistive devices was found in other studies. For example, in a recent study, Nucklos et al., showed that the optimal ankle stiffness from the assistive device should not be more than (around) 50% of the quasi stiffness of the ankle joint [77] which could be found by neuromuscular models.

**3) Feedback signal:** The last novelty of this study is introducing the ground reaction force (GRF) as a useful feedback signal. Basically, our control approach is similar to impedance control while the actuator impedance (stiffness) is adjusted by the GRF signal. Biological evidence supports implementation of GRF for locomotion control [40, 49]. Studies on locomotor disorders confirm the same concept of force feedback importance from another perspective when the proprioceptive inputs can adapt the locomotor pattern to external demands [78]. Dietz et al, stated that “cyclical leg movements <u>ONLY</u> in combination with loading of the legs lead to an appropriate leg muscle activation” [78]. In other words, the compensatory leg muscle activation during the stance phase of the gait is load dependent [41, 49]. Pathological gait analyses confirm the importance of GRF in locomotion control; e.g., when there is a reduced load sensitivity and respectively, decreased leg extensor activation in Parkinsonian gaits [39]. Inspired by these studies, positive force feedback (PFF) in muscle control [59, 63] and the FMCH template model are all in line with the idea of using GRF as a feedback signal for locomotion control. Although, control theory says that positive feedback destabilizes controlled systems, the PFF explains how reinforcing the extensor activity during the loading period of the stance phase will contribute to load compensation without leading to instability [49]. It is noteworthy to mention that in PFF, the constraints (limited muscle force) and hybrid dynamics of the gait (switching between stance and swing phases) are of reasons to explain how positive feedback could stabilize locomotion. In contrary to phase-detection-based methods (for gait assistance), here, the GRF-based control indirectly synchronizes the exosuit control with human movement. The GRF is considered as an informative sensory signal for control which removes the need to realize gait phasing. In equations (5) to (7) we removed the dependency to time, by using the GRF and the artificial muscle properties (e.g., length and force). In addition, external perturbations will be reflected in the GRF patterns and in the case of designing an appropriate controller, this sensory information could potentially robustify the system against perturbations. Technical issues exist in the implementation level such as measuring the GRF in a portable exosuit. Insole force sensor would be our solution to provide the GRF for overground walking in the future.

Comparison between the efficiency of the ABC and the optimal solution supports advantages of our bioinspired design. Although the ABC results in less metabolic cost reduction, it is more efficient than the optimal solution even if it is equipped with serial compliance. More interestingly, the developed device could generate efficiency above 100%. This phenomenon is also observed when passive exosuits can yield metabolic cost reductions without consuming energy [79–81]. The total energy of the human body and the assistive device can be decreased to less than the metabolic cost in the unassisted gait because of: 1) storing and returning energy with compliant elements [79–81] and 2) transferring energy from one joint to another with biarticular mechanisms [80, 81]. This is not investigated sufficiently in active devices due to influence of the actuators efficiency. Here, we used the mechanical efficiency which neglects the efficiency of the electric motors to focus on the control and mechanical design of the BAExo. We will further investigate that topic in our future studies on human assisted walking experiments.

## Supporting information

**S1 Video. The pilot experiment of the BAExo for the walking task.** The attached video shows the pilot experiment of the functionality of the BAExo assisting a subject in walking on an instrumented treadmill.

## Acknowledgments

We are thankful for the constructive discussions with Andre Seyfarth on the description of the experimental results. We also would like to thank Jennifer Nutter for her great support in revising the paper.

## Footnotes

1 For more information about control of assistive devices see [18, 19].

